# Leaf and root microbiome signatures of gray mangrove trees in the Red Sea

**DOI:** 10.64898/2026.09.22.753136

**Authors:** Diego J. Jiménez, Tahira Jamil, María Fernanda Peña-Valencia, Hanin Alzubaidy, Azad Baazeem, Kaitlyn O’Toole, Lucas William Mendes, Susana Carvalho, Alexandre S. Rosado

## Abstract

Mangroves persist under strong environmental constraints with support from microbes. Although microbial communities differ among mangrove compartments, their assembly mechanisms and compartment-specific functional signatures remain poorly resolved. Here, we combine peptide nucleic acid–clamping coupled with 16S rRNA gene amplicon sequencing and genome-resolved metagenomics to investigate the microbial communities associated with the leaves and roots of gray mangrove trees in the Red Sea. Our results suggest homogeneous selection and homogenizing dispersal as key processes of microbial community assembly in leaves. This phyllosphere hosts novel prokaryotic lineages, with some of its members probably synthesizing rhodopsins, plant polysaccharide–degrading enzymes, and γ-aminobutyric acid, a metabolite that increase tolerance to salinity stress. Genomes affiliated with *Desulfobacterales* and *Sedimenticolaceae* taxa are abundant belowground, suggesting complementary sulfur- and nitrogen-cycling capacities similar as occur in other blue-carbon ecosystems. Comparison with root-derived genomes from cordgrass revealed a host-driven selection of *Sedimenticolaceae* species with convergent metabolic profiles. Overall, this study provides an integrative view about the microbial biology of gray mangrove trees, offering foundational insights into the diversity, ecology, and predictive functionality of their aboveground–belowground microbiomes.

## Introduction

Mangroves occupy the dynamic interface between land and sea and provide globally important ecosystem services, including coastal protection, nursery habitats for a wide range of organisms (e.g., fishes, bivalves, crabs, insects, and microbes), carbon storage, and climate regulation (Alongi, 2014; Trevathan-Tackett et al. 2019; Getzner et al. 2020). Their persistence under harsh and fluctuating environmental conditions (e.g., high salinity, intense solar radiation, nutrient limitation, and/or anoxia) depends not only on plant physiology but also on their associated microorganisms. In fact, microbial communities inhabiting mangrove trees (e.g., leaves and roots) and the surrounding sediments contribute to nutrient cycling (including carbon, nitrogen, sulfur, and phosphorus) and organic matter turnover, along with enhancing the tolerance of mangroves to biotic and abiotic stressors (Wainwright et al. 2023; Ding et al. 2025; Huang et al. 2025). These mangrove-associated microbiomes are considered key components for improving mangrove restoration and rehabilitation strategies (Allard et al. 2020; Ohimain et al. 2026).

For decades, microbial communities associated with mangrove soils and sediments have been well-described and characterized (Andreote et al. 2012; Jiménez et al. 2015; Alzubaidy et al. 2016; Purahong et al. 2019; Qian et al. 2023; Liu et al. 2025; Sidharthan et al. 2025; Jiménez et al. 2025). Furthermore, advances in molecular methods, declining sequencing expenses, and innovative bioinformatic tools have facilitated the analysis of microbial profiles across different mangrove compartments (e.g., leaves, steams, roots, and the rhizosphere) (Zhuang et al. 2020; Sui et al. 2023; Yang et al. 2023; Yuan et al. 2023; Hsiao et al. 2024). For instance, a mangrove “core” microbiome was proposed after analyzing 16S rRNA gene sequencing data obtained from leaves, fruits, roots, and sediment samples associated with *Avicennia alba* and *Sonneratia alba*, two common mangrove species in Southeast Asia (Wainwright et al. 2023). A more recent study characterized leaf- and root-associated microbiomes in *A. germinans* and *Rhizophora mangle* (from French Guiana) using 16S rRNA gene amplicon sequencing (Vigneron et al. 2026). Another two studies using similar approaches also described microbial communities associated with *A. marina* (gray mangrove), one of the most abundant mangrove species in the Red Sea (Ghabban et al. 2024; Alghamdi et al. 2024).

When characterizing plant-associated microbiomes using 16S or 18S rRNA gene amplicon sequencing, the coextraction of host-derived DNA remains a common technical challenge, which can result in the unintended amplification of chloroplast and mitochondrial rRNA gene sequences (Víquez-R et al. 2020; Taerum et al. 2020). This problem can drastically reduce the counts of microbial-derived sequences, obstructing the detection of low-abundant taxa and biasing microbial diversity profiles (Hussain et al. 2025). Fortunately, specific peptide nucleic acid (PNA) clamps provide a practical solution for minimizing the coamplification of host DNA, thus improving microbial resolution in plant-associated microbiome studies (Fitzpatrick et al. 2018). PNAs are artificially synthesized oligomers that bind to specific host sequences, thereby blocking their amplification during PCR (Taerum et al. 2020). Although PNA-clamping has been recently applied in studies of maize, wheat, and oak microbiomes (Mukhtar et al. 2025; Dubois et al. 2025; Hussain et al. 2025), its potential for improving resolution in mangrove-associated microbiome surveys—particularly in compartments such as leaves and roots—remains underexplored (Yuan et al. 2023; Alghamdi et al. 2024; Vigneron et al. 2026).

In mangrove-associated soil and sediments, the analysis of metagenome sequences and the reconstruction of gene and genome catalogs have enabled researchers to clarify the ecological roles and metabolic potential of abundant microbial taxa (Zhang et al. 2023; Liu et al. 2025; Bohra et al. 2025; Peña-Valencia et al. 2026; Balvino-Olvera et al. 2026). Furthermore, microbiomes associated with other coastal/marine plants, such as seagrass and saltmarsh cordgrass (*Spartina alterniflora*), have been characterized using genome-resolved metagenomics (Liu et al. 2023; Sun et al. 2024; Sun et al. 2026; Rolando et al. 2024). These studies have linked microbial genomes and taxa to specific functions and traits. Conversely, mangrove tree compartments remain comparatively underexplored, and metagenomic data obtained from leaves and roots are still scarce, despite recent studies in *A. marina* (Bohra et al. 2025) and *A. germinans* (Lemos-Lucumi et al. 2025). These studies have reported that leaf-associated microbial communities may contribute to amino acid metabolism, while root-associated microbial communities may help to mitigate plant stress and perform nitrate- and sulfur-reduction processes. Nevertheless, the lack of comprehensive metagenomic analyses in mangrove leaves and roots impairs a deeper understanding of their microbial signatures (Krummenauer et al. 2025; Sujeeth et al. 2026), which ultimately support the growth, health, and stress resilience of these ecosystems (Trevathan-Tackett et al. 2019).

In the Red Sea, *A. marina* trees persist under harsh environmental conditions that impose markedly different constraints on aboveground and belowground compartments. In this scenario, leaves and roots represent fundamentally distinct microbial habitats. Leaves are exposed to intense solar radiation and extreme salinity, whereas roots may experience low oxygen levels, nutrient limitation, sulfide toxicity, and pronounced chemical gradients. We hypothesized that these contrasting environmental filters shape microbial community structure and assembly processes, favoring distinct functional strategies in both mangrove tree compartments. Such microbial signature partitioning may represent a common organizing principle in other blue-carbon plants. To test this hypothesis, we (i) compared the microbial communities associated with the leaves, roots, rhizosphere, and sediments of *A. marina* trees in the northern Red Sea, evaluating the effect of the PNA-clamping method on the prokaryotic diversity profiles on leaf and root samples, and (ii) investigated the microbial signatures of *A. marina*-derived roots and leaves via genome-resolved metagenomics, comparing them with those found in saltmarsh cordgrass–associated microbiomes. By integrating cutting-edge molecular approaches, community ecology, and genome-centric comparative analyses, this study moves beyond describing the gray mangrove microbiome to unraveling how microbial signatures are structured across blue-carbon plant compartments and how they may support the persistence of mangrove trees in the Red Sea.

## Results

### Contrasting microbial profiles between gray mangrove-associated compartments

A representative sampling of *A. marina* leaves (n = 33), roots (n = 33), rhizosphere (n = 16), and sediments (n = 46) was conducted across at six different locations in the northern Red Sea (Fig. 1A). To compare among compartments, we used mangrove leaf and root samples subjected to the PNA-clamping process (see next section). After removing chloroplast- and mitochondrial-like sequences, 9656 prokaryotic-derived amplicon sequence variants (ASVs) were obtained in the 16S rRNA gene dataset. As anticipated, each plant compartment and sediment samples contained a particular microbial community (Fig. 1B), with significant differences being observed between them (permutational multivariate analysis of variance (PERMANOVA); adonis; F = 5.601; R^2^ = 0.157; P < 0.001). Leaf-associated microbial communities were highly dissimilar compared with other mangrove-associated microbiomes, showing low and significant (Kruskal– Wallis; P < 0.001) alpha diversity values (i.e., Shannon index and Observed species) (Fig. 1C). Taxonomic plots at order level revealed that, on average, leaf-associated microbial communities were dominated by *Rhodobacterales*, *Halobacteriales* (*Archaea*), and *Salinisphaerales*, whereas roots, rhizosphere, and sediments contained high relative abundances of ASVs affiliated with *Desulfobacterales*, *Hyphomicrobiales*, and *Actinomarinales*. Furthermore, seawater (n = 4) surrounding the mangrove trees was primarily dominated by *Flavobacteriales*, *Rhodobacterales*, and *Synechococcales* (Fig. 1D).

**Figure 1.**
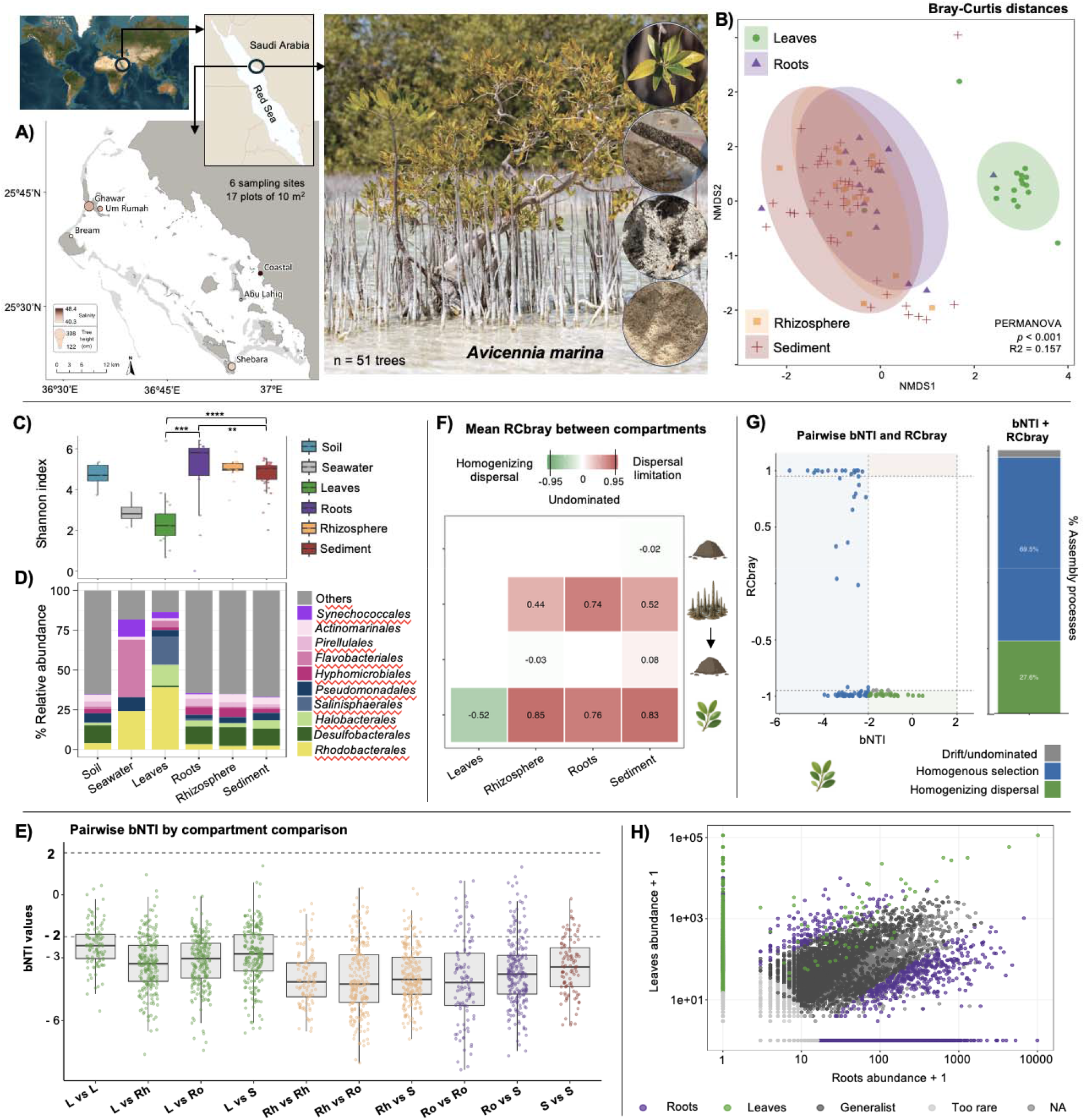
Prokaryotic diversity analyses in *A. marina*-associated microbiomes via 16S rRNA gene amplicon sequencing. **A)** Sampling locations in the Al Wajh Lagoon in the northern Red Sea (Saudi Arabia) and representative pictures of *A. marina* tree, mangrove compartments (i.e., leaves, roots, and rhizosphere), and sediment samples (from top to bottom). **B)** Nonmetric multidimensional scaling (NMDS) plot depicting Bray–curtis distances among leaf, root-, rhizosphere-, and sediment-associated microbial communities. **C)** Comparison of Shannon index values among leaf-, root-, rhizosphere-, and sediment-associated microbial communities, including controls (i.e., surrounding seawater and soil without mangrove vegetation). Significant differences were found among leaves, roots, and sediments (ANOVA; ** P < 0.05 and **** P < 0.0001). **D)** Taxonomic plot at order level depicting the top 10 abundant taxa in each *A. marina*-associated microbiome and control. **E)** Distribution of pairwise βNTI values among and within sediment (S), rhizosphere (Rh), root (Ro), and leaf (L) samples. Dashed horizontal lines indicate βNTI thresholds of −2 and +2. Values <−2 indicate homogeneous selection, values >+2 indicate variable selection, and values between −2 and +2 indicate stochastic or undominated assembly. **F)** Mean Bray–Curtis-based Raup–Crick metric (RC_bray_) values among pairwise comparisons. Positive values indicate communities that are more dissimilar than expected under the null model, whereas negative values indicate communities that are more similar than expected. **G)** Pairwise relationship between βNTI and RC_bray_ values for leaf samples. Dashed vertical lines indicate βNTI thresholds of −2 and +2, and dashed horizontal lines indicate RC_bray_ thresholds of −0.95 and +0.95. Points represent pairwise comparisons among leaf samples and are colored according to the integrated assembly process classification. **H)** Pairwise CLAM classification. The panel illustrates a comparison between root and leaf microbial communities. Points represent individual ASVs classified as specialists of one treatment, generalists shared between treatments, or extremely rare to classify. Specialist ASVs are colored according to their associated treatment, whereas generalists and rare ASVs are depicted in gray. Axes represent total ASV abundance in each treatment plus one, plotted on a log10 scale.

Based on the co-occurrence networks of ASVs, root-associated microbiomes may be the most complex system with 981 nodes and 18,402 edges. The beta-nearest taxon index (βNTI) indicated a predominance of homogeneous selection as a major process of community assembly (βNTI <−2) in all gray mangrove compartments (Fig. 1E). Furthermore, a taxonomic null model inference using the mean Bray–Curtis-based Raup–Crick metric (RC_bray_ values) indicated that microbial community turnover from all compartments was governed by ecological drift, weak selection, or multiple nondominant processes (Fig. 1F). Root-associated microbial communities demonstrated the strongest evidence of divergent assembly or dispersal limitation, whereas leaf-associated microbial communities could be shaped by environmental filtering and homogenizing dispersal (Figs. 1F and G). Moreover, using a multinomial species classification method (CLAM), we found that root-associated microbial communities demonstrated the strongest specialization signal, where 1111 and 3631 ASVs were classified as specialists compared with the rhizosphere- and leaf-associated datasets, respectively (Fig. 1H).

### Enhancing diversity resolution in gray mangrove-associated microbiomes

The impact of PNA-clamping on prokaryotic diversity profiles was assessed in a pair-set of leaf (n = 17) and root (n = 15) samples. Briefly, our results revealed that PNA-clamping reduced the relative abundance of sequences affiliated to chloroplasts, in turn increasing the amount of sequencing data from *Bacteria* and *Archaea* (Fig. 2A). In leaf samples, ∼90% of raw 16S rRNA gene sequences belonged to chloroplast when PNA-clamping was not used, and this proportion decreased to ∼25% when PNA-clamping was applied (Fig. 2A). As an effect of the depletion of chloroplast sequences, a higher microbial community variability was detected in leaf and root samples. Nevertheless, the use of PNA-clamping exerted no effect on the beta diversity profiles of leaf-associated (PERMANOVA; adonis; F = 2; R^2^ = 0.061; P > 0.05) and root-associated (PERMANOVA; adonis; F = 1.4; R^2^ = 0.045; P > 0.06) microbial communities (Fig. 2B). Furthermore, Shannon index values increased significantly when PNA-clamping was used, especially in root samples (Kruskal–Wallis; P < 0.0001) (Fig. 2C).

**Figure 2.**
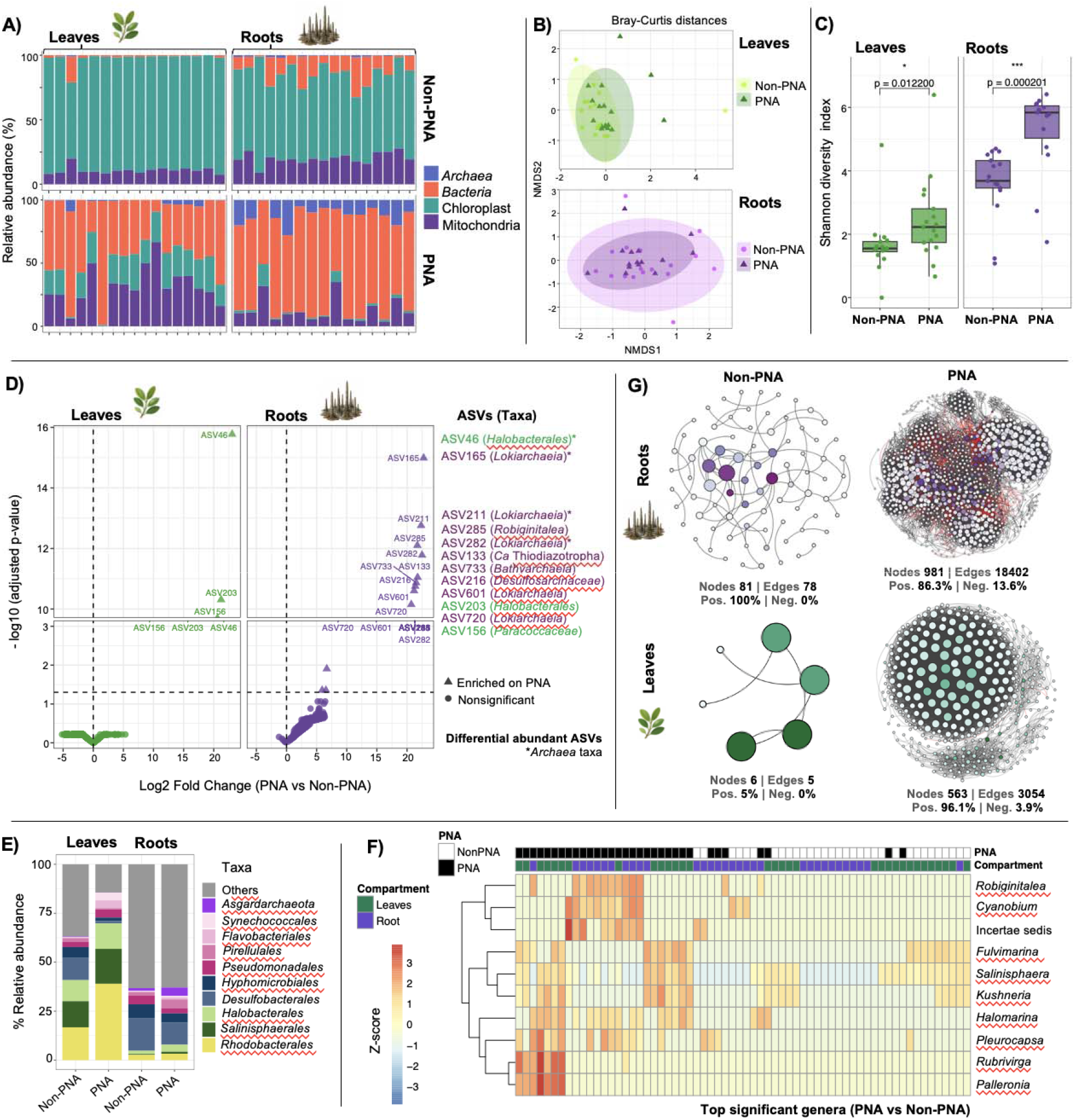
Effect of peptide nucleic acid (PNA)-clamping on the prokaryotic diversity of gray mangrove leaf- and root-associated microbiomes. **A)** Comparison of 16S rRNA gene sequencing raw data obtained from leaves and roots with and without PNA-clamping. The plot depicts the proportion of sequences affiliated to *Bacteria*, *Archaea*, Chloroplast, and Mitochondria. **B)** Nonmetric multidimensional scaling (NMDS) plot depicting Bray–Curtis distances between leaf and root samples with and without PNA-clamping. No statistical differences (permutational multivariate analysis of variance (PERMANOVA); P > 0.01) were found between PNA- and non-PNA-treated samples. **C)** Shannon diversity index values of leaf-and root-derived samples with and without PNA-clamping. **D)** Differential abundance analysis via DESeq2. A volcano plot is depicted to identify highly enriched ASVs when PNA-clamping was used in both plant compartments (leaf-derived ASVs in green and root-derived ASVs in purple). **E)** Taxonomic plots at order level displaying the top 10 abundant taxa, comparing samples treated with PNA or untreated with PNA in leaves and roots. **F)** Heatmap representing the normalized relative abundance of the top 10 genera that were significantly enriched (DESeq2) in PNA- and non-PNA-treated leaf and root samples. **G)** Prokaryotic co-occurrence networks inferred from the SparCC analysis. The SparCC correlations were based on a magnitude of >0.7 (positive correlation) or <−0.7 (negative correlation). Purple and green nodes represent highly connected ASVs, and the size of the nodes is proportional to the number of connections (degree).

A differentially abundant analysis revealed significant enrichment (P < 0.001; log2-fold change > 15) of three ASVs on leaf samples treated with PNA versus non-PNA. These ASVs were affiliated with *Paracoccaceae* and *Halobacteriales* (Fig. 2D). In roots, nine ASVs exhibited significantly (P < 0.001; log2-fold change > 15) higher relative abundance on PNA-treated samples compared with that on non-PNA-treated samples. Among the nine ASVs, six were affiliated with *Archaea* species (*Lokiarchaeia* and *Bathyarchaeia*), and three with bacterial taxa (*Robiginitalea*, *Ca.* Thiodiazotropha, and *Desulfosarcinaceae*) (Fig. 2D). Leaf-derived samples subjected to PNA-clamping showed a higher abundance of *Rhodobacterales* of ∼40% compared with samples not subjected to PNA-clamping (Fig. 2E). Moreover, ASVs affiliated with *Kushneria* and *Salinisphaera* were considerably abundant on PNA-treated samples compared with that on root samples (Fig. 2F). The ASV co-occurrence network analyses revealed that using PNA-clamping in leaf and root samples considerably increased the number of edges and nodes (Fig. 2G), suggesting an enhancement in the assessment and resolution of prokaryotic diversity profiles.

### Taxonomic affiliation of high-quality metagenomic reads

Metagenomic sequencing from gray mangrove leaf (n = 18) and root (n = 20) samples yielded approximately 169 and 161 Gbp of sequence information, respectively. High-quality sequences were taxonomically classified using Kraken/Bracken and the PlusPFP database, resulting in ∼60% of short reads unclassified at kingdom level in leaf samples, most probably belonging to prokaryotes. A small fraction of the total sequences in leaves was affiliated to *Archaea,* primarily from the order *Halobacteriales* (Fig. 3A). In root samples, there was an increase in the proportion of reads affiliated to *Bacteria* with a decrease in the proportion of plant-derived reads compared with that in leaf samples.

**Figure 3.**
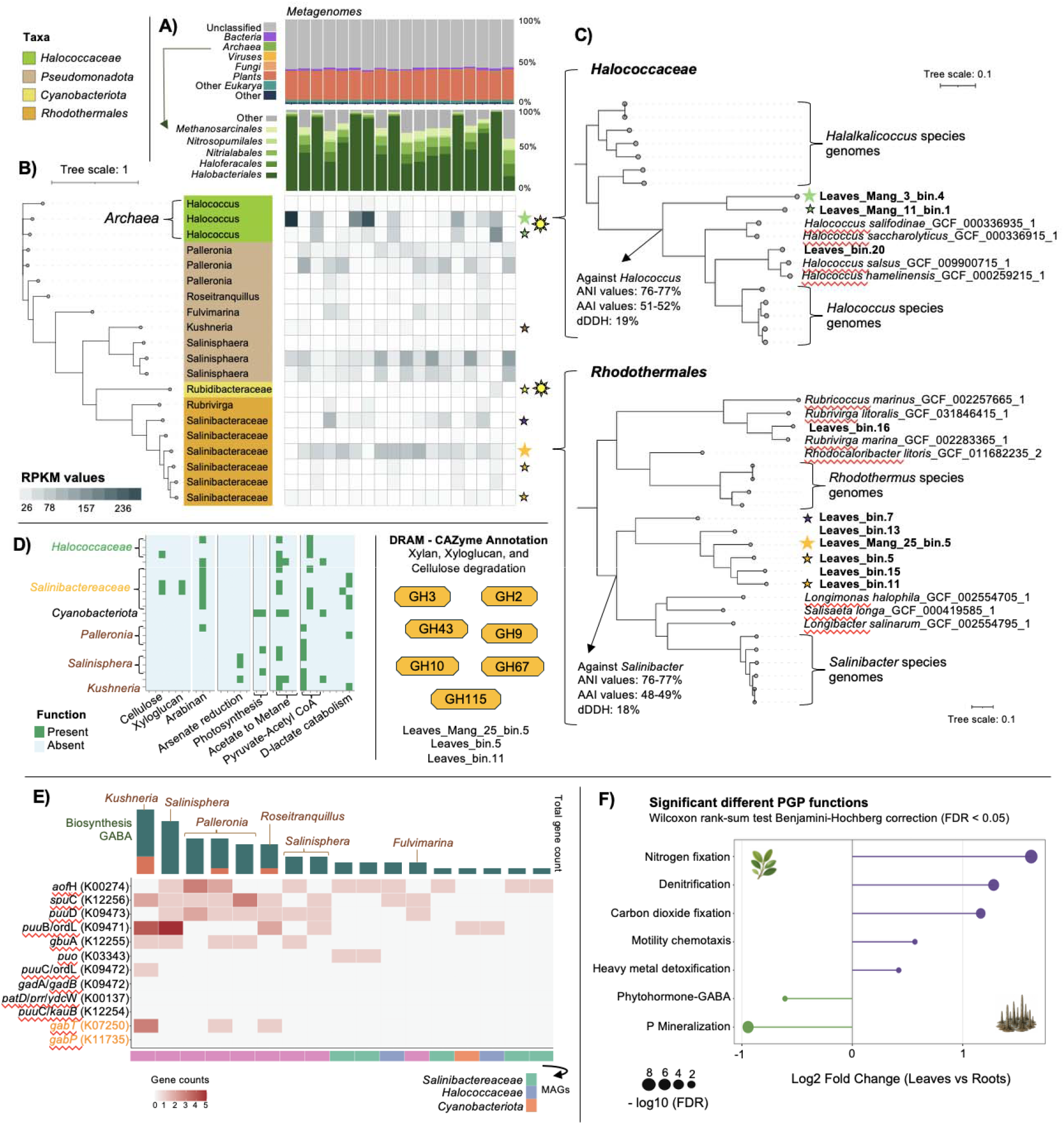
Genome-resolved metagenomic analyses in gray mangrove leaf-derived samples. **A)** Read-based taxonomic affiliation of metagenomic sequences using the Kraken– Braken software. The bar plot in purple shows the five most abundant orders within *Archaea*-assigned clean sequences. **B)** Phylogenomic tree (built using GToTree v16.31 with a prepacked single-copy gene set) of 20 dereplicated medium-to-high-quality MAGs obtained after the coassembly of 18 metagenome datasets. The phylogenomic tree illustrates taxonomic affiliation based on GTDB-tk (left), and the reads per kilobase per million mapped reads (RPKMs) values, obtained after mapping back unassembled sequences, are plotted in the heatmap. Colored stars represent high-quality MAGs (<5% contamination and >90% completeness) within the dataset, and sun icons represent MAGs with the presence of photorhodopsins (K04641 and/or K04642). **C)** Two distinct phylogenomic trees constructed using type representative genomes from the *Halococcaceae* family (top) and *Rhodothermales* order (bottom), which were identified via MiGA using TypeMat database release 2025-08, and MAGs from our dataset (in bold). **D)** Functional gene annotation (using the DRAM pipeline) of MAGs derived from gray mangrove leaf samples. The heatmap depicts functions associated with the presence/absence of specific carbohydrate-active enzyme (CAZy) families or KEGG orthology groups predicted to be involved in different metabolic processes. CAZy families found in *Salinibacteraceae*-affiliated MAGs and involved in the degradation of xylan, xyloglucan, and cellulose are depicted in yellow. **E)** Gene-level characterization of γ-aminobutyric acid (GABA) metabolism in leaf-associated MAGs. Heatmap depicting the distribution of gene counts assigned to GABA biosynthesis (green) and conversion (orange) across MAGs; color intensity represents gene count. Phylum-level taxonomic affiliation of MAGs is displayed at the bottom of the heatmap. *Pseudomonadota*-affiliated MAGs are highlighted at the top of the figure. **F)** Plot illustrating fold change values of significant functions (plant-growth-promoting traits) between MAGs derived from leaves and roots.

### Metagenome-assembled genomes from gray mangrove leaves

After the coassembly of metagenomic sequences (N50 of 57,681 bp) and binning of 91,870 contigs (⋝1000 bp), 51 medium-to-high-quality (≥50% completeness and <10% contamination) metagenome-assembled genomes (MAGs) were obtained. The size of 20 dereplicated (at 95% average nucleotide identity (ANI)) leaf-derived MAGs ranged from 1.6 to 4.1 Mbp. Based on GTDB-tk, six MAGs probably belonged to species of the family *Salinibacteraceae*. Furthermore, *Halococcus*, *Salinisphaera*, and *Palleronia* were represented by three MAGs each. Single MAGs obtained from *Fulvimarina*, *Kushneria*, *Rubrivirga*, and a *Cyanobacteria* species were also detected in the dereplicated dataset (Fig. 3B). These results suggest an acceptable representation of the gray mangrove leaf microbiome compared with the 16S rRNA–based taxa composition.

### Unveiling hidden microbial taxa in the gray mangrove phyllosphere

The RPKM (reads per kilobase per million mapped reads) values indicated that two high-quality MAGs (Leaves_Mang_11_bin.1 and Leaves_Mang_3_bin.4) affiliated with *Halococcus* species were consistently abundant in most leaf samples (Fig. 3B). These two MAGs exhibited lower values of ANI (<78% in all pairwise comparisons) and digital DNA–DNA hybridization (dDDH) (<19%) with genomes from *Halococcus* species (Fig. 3C). Both MAGs also demonstrated ∼52% average amino acid identity (AAI) with *Halococcus thailandensis* (GCF_000336715.1). A phylogenetic comparison of these two MAGs with representative genomes from *Halococcaceae* suggested that they belong to a novel genus (the proposed name is *Halofoliata photomarina* gen. nov. sp. nov.) within this family (P = 0.42 based on Microbial Genome Atlas (MiGA)). This assumption was highly supported by the 16S rRNA gene phylogenetic tree constructed using information obtained from bacterial-type strains.

A high-quality MAG (Leaves_Mang_25_bin.5), most probably belonging to the order *Rhodothermales* (P = 0.33 based on MiGA), was consistently abundant in leaf samples (Fig. 3B). Based on its low ANI (∼77%), AAI (∼49%), and dDDH (∼18%) values compared with those of representative genomes from *Salinibacter* species, we suggest that this MAG belongs to a novel bacterial genus (the proposed name is *Salinimangrovia arabica* gen. nov. sp. nov). This claim was supported by a phylogenomic tree where a set of three high-quality MAGs, including Leaves_Mang_25_bin.5, was grouped separately compared with representative genomes from *Rhodothermales* (Fig. 3C). Furthermore, Leaves_bin.7 may belong to another genus within this order, exhibiting 63.9% of AAI compared with Leaves_Mang_25_bin.5. Unfortunately, 16S rRNA gene sequences were not recovered from these four MAGs. Nevertheless, a phylogenomic tree, based on Genome BLAST Distance Phylogeny (GBDP) distances, supports their novelty compared with other bacterial-type strains.

A single high-quality MAG affiliated to a *Rubidibacteraceae* species (Leaves_bin.17) may also represent a novel cyanobacterial taxon. It exhibited an AAI value of 53.9% with *Lusitaniella coriacea* (GCF_050053475.1); however, it was not further investigated owing to uniqueness and low abundance compared with other MAGs. Moreover, all the prokaryotic species and genus names proposed in this study have been submitted to the SeqCode registry.

### Genomic-based physiological traits of novel prokaryotic lineages

We predicted the physiological traits of the two proposed novel taxa using type MAGs (Leaves_Mang_3_bin.4 and Leaves_Mang_25_bin.5) and the metaTraits software. Briefly, our results demonstrated that both species may grow in the presence of oxygen, at pH 7.4–7.8, and temperature 30°C–42°C. The optimum salinity for *Halofoliata* spp. is ∼16% NaCl, whereas it is 8% NaCl for *Salinimangrovia* spp. Both species may utilize D-xylose, D-mannose, citrate, and hydrolyzed starch. Nevertheless, only *Salinimangrovia* species are predicted to utilize D-mannitol, L-arabinose, and lactose. It may also hydrolyze casein and convert nitrate into nitrite.

### Predictive functions of abundant species in the gray mangrove phyllosphere

Functional annotation of MAG-derived genes was conducted using three computational pipelines, viz., DRAM, PGPg_finder, and eggNOG-mapper (see methods). Genes encoding bacteriorhodopsin (K04641) and halorhodopsin (K04642) were detected in MAGs affiliated with the proposed novel *Halofoliata* species. These genes were absent in Leaves_bin.20 (affiliated with *Halococcus*). The bacteriorhodopsin gene (K04641) was found in Leaves_bin.17, a MAG affiliated with a *Cyanobacteria* species (Fig. 3B). This cyanobacterial gene exhibited 63.5% amino acid similarity (E-value 1e-68) with a membrane protein of *Chroococcidiopsis* sp. (WP_250123691.1) and contained a YesE (COG3631) domain. Moreover, MAGs affiliated to *Halofoliata* species contained genes encoding V/A-type ATPases, proteins involved in ATP synthesis via the pumping of sodium ions across the cell membrane.

Three MAGs, which clustered together with the proposed *Salinimangrovia arabica* gen. nov. sp. nov., contained genes encoding glycosyl hydrolases families (e.g., GH3, GH9, GH10, and GH43) that are potentially involved in the degradation of plant-derived polymers such as cellulose, xyloglucan, and arabinan (Fig. 3D). These three MAGs also harbored genes potentially involved in phosphorus (P) mineralization and acetate and lactate catabolism (Fig. 3D; Fig. 4A). MAGs belonged to *Salinisphaera* contained genes potentially involved in arsenate reduction (Fig. 3D). Interestingly, MAGs belonged to *Salinisphaera*, *Palleronia*, and *Kushneria* contained genes (e.g., *puu*BCD, *spu*C, *aof*H, and *gbu*A) predicted to be involved in the biosynthesis of γ-aminobutyric acid (GABA) primarily via the Puu (putrescine utilization) pathway (Fig. 3E). Genes predicted to confer these two plant-growth-promoting (PGP) traits (i.e., GABA production and P mineralization) were significantly (Wilcoxon test; Benjamini– Hochberg correction; FDR < 0.05) present in MAGs derived from leaves compared with that in MAGs obtained from roots (Fig. 3F; Fig. 4A).

**Figure 4.**
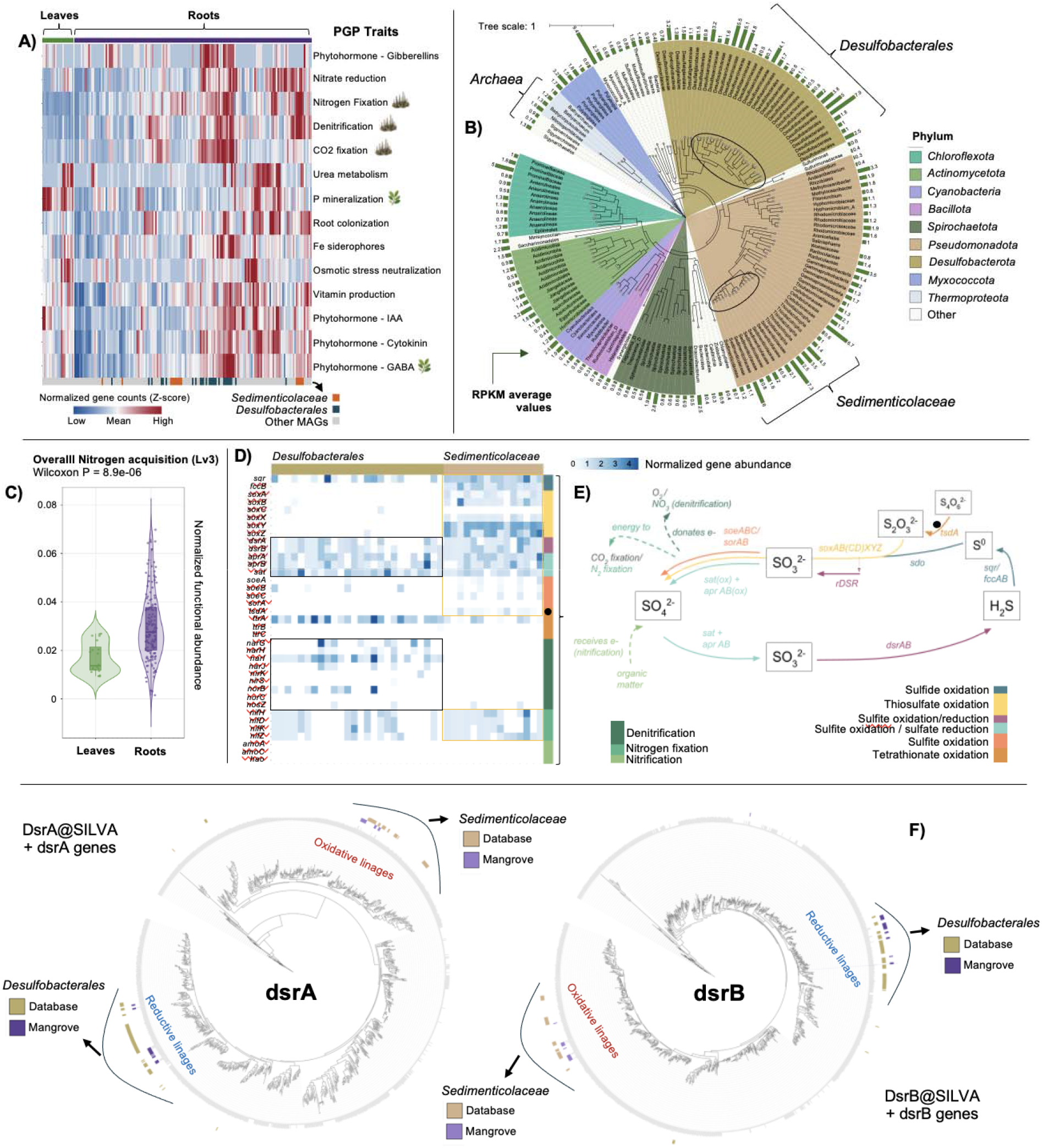
Functional annotation and taxonomy classification of metagenome-assembled genomes obtained from gray mangrove leaves and roots. **A)** Functional annotation of MAG-derived genes using PGPg_finder. The heatmap depicts normalized gene counts affiliated to different PGP traits across MAGs derived from leaves and roots. At the bottom of the heatmap, MAGs affiliated to *Desulfobacterales* (blue) and *Sedimenticolaceae* (orange) are highlighted. Icons of leaves and roots represent traits significantly enriched in each compartment. **B)** Phylogenomic tree of the dereplicated set of MAGs with medium-to-high quality obtained from 20 root samples. The tree was constructed using GToTree v16.31 with a prepacked single-copy gene set. Taxonomy was determined using GTDB-tk. Outside the tree, the average reads per kilobase per million mapped reads (RPKMs) values per MAG are depicted. MAGs affiliated with *Archaea* phylum, *Desulfobacterales* order, and *Sedimenticolaceae* family are highlighted with brackets. **C)** Normalized abundance of genes involved in nitrogen acquisition (Level 3 on PGPT ontology) found in leaf- and root-derived MAGs. **D)** Heatmap depicting gene count values corrected by MAG completeness (rows) involved in nitrogen and sulfur transformation process across root-derived MAGs (columns) affiliated to *Desulfobacterales* order and *Sedimenticolaceae* family. **E)** Schematic of the major pathways and genes involved in the sulfur cycle coupled with the major process in the nitrogen and carbon cycle. **F)** Phylogenetic tree of DsrA and DsrB genes found in MAGs affiliated to *Desulfobacterales* and *Sedimenticolaceae* and genes deposited in the dsrAB@SILVA database. *Sedimenticolaceae-*derived genes clustered with oxidative bacterial lineages, whereas *Desulfobacterales*-derived genes grouped with reductive bacterial lineages.

### Sulfur and nitrogen reducers/oxidizers found in gray mangrove roots

After the coassembly of metagenome sequences (N50 of 3135 bp) and binning of 8,96,843 contigs (⋝1000 bp), 222 medium-to-high-quality MAGs were reconstructed. The sizes of 153 dereplicated sets of root-derived MAGs were in the range of 0.5–9.8 Mbp. Most of these MAGs belong to *Pseudomonadota* (n = 41), *Desulfobacterota* (n = 33), and *Actinomycetota* (n = 14) phyla. In addition, seven MAGs were affiliated with *Archaea* species (Fig. 4B). Based on RPKMs values, 26 and 15 MAGs belonged to the order *Desulfobacterales* and the family *Sedimenticolaceae*, respectively, were abundant across the root samples. Among them, six MAGs were affiliated with *Ca.* Thiodiazotropha. Moreover, specific MAGs affiliated with other bacterial taxa, e.g., *Myxococcota*, *Rhizobiales*, *Rariloculaceae*, *Cellvibrionaceae*, and *Promineifilaceae*, exhibited also high RPKM values (Fig. 4B).

Compared with leaf-derived MAGs, genomes obtained from root samples contained significantly (Wilcoxon test; Benjamini–Hochberg correction; FDR < 0.05) more genes involved in nitrogen fixation, denitrification, and carbon fixation (Fig. 4A; Fig. 3F). The normalized proportion of genes potentially involved in nitrogen acquisition was considerably higher in root-derived MAGs than in leaf-derived MAGs (Fig. 4C). Functional gene annotation of root-derived MAGs using DRAM, PGPg_finder, and eggNOG-mapper suggested that some of them affiliated to the family *Sedimenticolaceae* have the potential for fixing nitrogen (harboring the genes *nif*HDKZ) and oxidizing sulfur; whereas *Desulfobacterales*-affiliated MAGs have the potential for reducing nitrogen and sulfur. Compared with *Sedimenticolaceae*-affiliated MAGs, *Desulfobacterales*-affiliated MAGs contained significantly more genes (Wilcoxon test; Benjamini–Hochberg correction; FDR < 0.05) involved in the production of phytohormones and organic volatiles, fixation of carbon dioxide, and detoxification of heavy metals.

*Sedimenticolaceae*-affiliated MAG contained genes involved in the oxidation of thiosulfate (*sox*ABXYZ), sulfite (*soe*ABC), tetrathionate (*tsd*A), and sulfide (*sqr* and *fcc*B). As complement, *Desulfobacterales*-affiliated MAGs contained genes involved in the reduction of sulfate (*sat* and *apr*AB) and sulfite (*dsr*AB) (Figs. 4D and E). The presence of *dsr*AB genes, together with other sulfur oxidation–related genes, in *Sedimenticolaceae-*affiliated MAGs suggests that they could operate in the oxidative direction (rDSR). To confirm this assumption, we compared both genes with reference sequences from the DsrAB@SILVA database described by Diao et al. (2023) and found that *Sedimenticolaceae* DsrA and DsrB sequences clustered with reference sequences belonging to the oxidative bacterial-type DsrAB lineage. Conversely, DsrA and DsrB sequences derived from *Desulfobacterales*-affiliated MAGs clustered with reference sequences of the reductive bacterial-type DsrAB lineages (Fig. 4F).

### *Sedimenticolaceae*: a signature in the radicular system of two blue-carbon plants

A comparative analysis of a dereplicated set of root-derived MAGs obtained from the gray mangrove (n = 153) (this study) and saltmarsh cordgrass (n = 164) (Huang and Pettersen et al. 2026) was conducted. Both coastal plant roots demonstrated an overrepresentation of MAGs affiliated with sulfate-reducing (e.g., *Desulfobacterales*) and sulfur-oxidizing (e.g., *Chromatiales*, *Campylobacterales*, and *Rhizobiales*) taxa (Figs. 5A–C). A phylogenomic tree revealed that MAGs derived from both ecosystems were found in almost all clades, although some of them were host-specific (Fig. 5A). Despite some similarities between both datasets, microbial communities are shaped by the host and MAGs in common (ANI > 95%) were not detected. This result was represented by a pairwise comparison between MAGs affiliated with the family *Sedimenticolaceae* in both datasets (Fig. 5B). Nevertheless, in an AAI pairwise comparison, three clusters (AAI > 70%) of MAGs were obtained, which contained representative genomes from cordgrass and gray mangrove roots (Fig. 5D).

**Figure 5.**
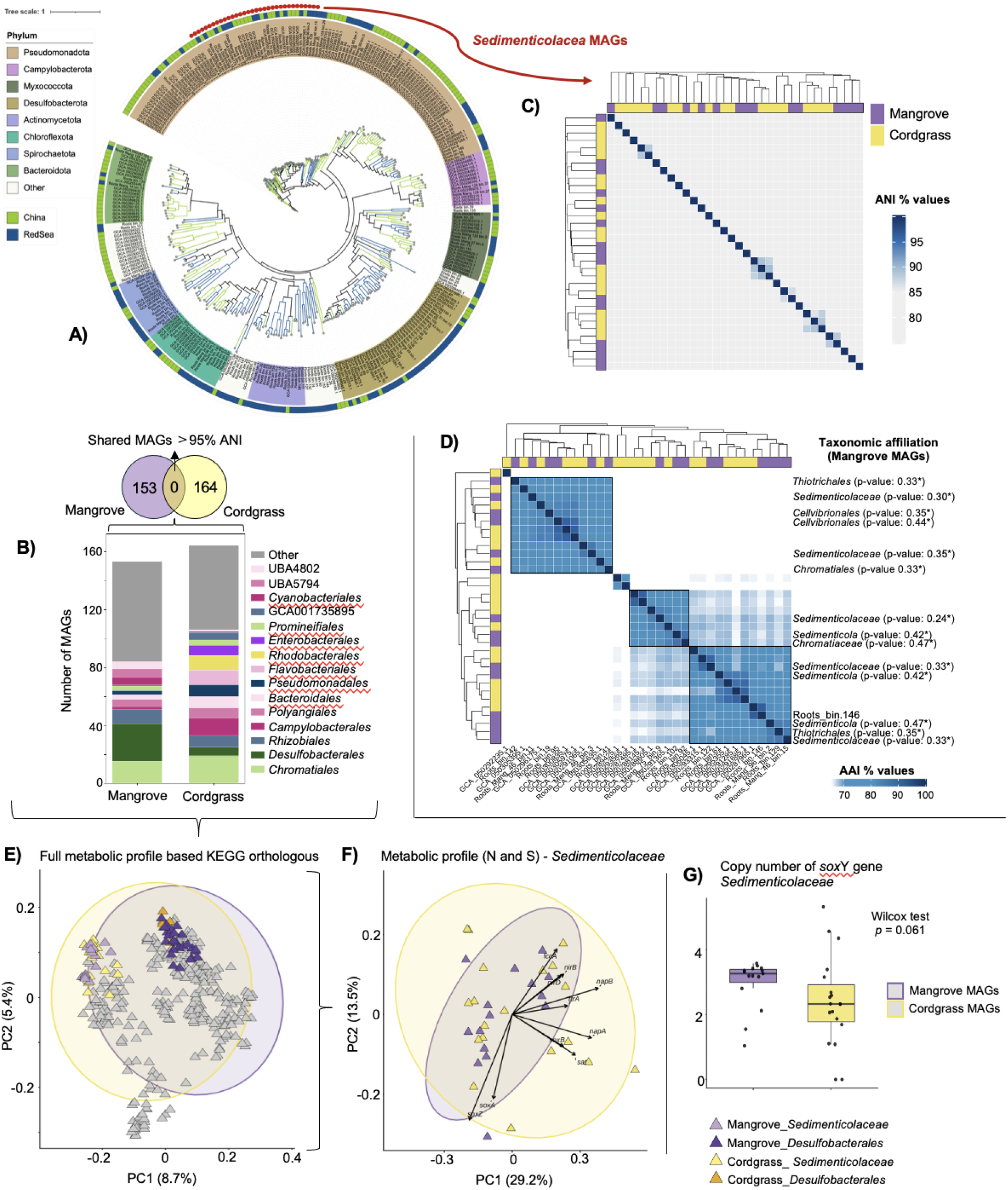
Comparison of metagenome-assembled genomes obtained from gray mangrove and saltmarsh cordgrass roots. **A)** Phylogenomic tree of a high-quality and dereplicated set of MAGs obtained from gray mangrove roots (green) and saltmarsh cordgrass roots (blue) (Huang and Pettersen et al. 2026). Red dots outside the tree represent MAGs affiliated to *Sedimenticolaceae.* **B)** Bar plot depicting the taxonomic affiliation of MAGs (via GTDB-tk) retrieved from mangrove (this study) and cordgrass roots. **C)** Average nucleotide identity (ANI) pairwise comparison between MAGs affiliated to *Sedimenticolaceae* in mangrove and cordgrass roots. **D)** Average amino acid identity (AAI) pairwise comparison between MAGs affiliated to *Sedimenticolaceae* in gray mangrove and cordgrass roots. Taxonomic affiliation, based on the Microbial Genomes Atlas (MiGA) webserver, of MAGs obtained from gray mangrove roots is illustrated at the right. **C)** Metabolic profile based on the presence/absence KEGG ortholog annotations. Clustering is based on Jaccard distances. MAGs belonging to *Desulfobacterales* and *Sedimenticolaceae* are highlighted with colors. **D)** Metabolic profile of MAGs affiliated to *Sedimenticolaceae* in both datasets using only completeness-corrected abundances of gene counts involved in nitrogen and sulfur transformations. Clustering is based on Bray–Curtis distances. **E)** Copy numbers of *soxY* (K17226) in MAGs affiliated to the *Sedimenticolaceae* family from gray mangrove and cordgrass datasets.

Regarding the complete functional profile determined using the presence/absence of KEGG orthologs (Fig. 5E), statistical differences (PERMANOVA, P < 0.001) were found between MAGs obtained from the gray mangrove versus MAGs obtained from cordgrass. However, the ecosystem source explained only 1.8% of the total variation (R^2^ = 0.018). Moreover, the significant PERMDISP result (P < 0.001) indicated high within-group variability, suggesting that the significant PERMANOVA result is affected by differences in dispersion rather than by a strong separation between both ecosystems. Furthermore, a clear separation was found in MAGs affiliated to *Desulfobacterales* and *Sedimenticolaceae*, which was independent of the ecosystem source and suggests that they are two different functional guilds (Fig. 5E).

A specific comparison between *Sedimenticolaceae*-affiliated MAGs obtained from both blue-carbon ecosystems suggested that they possess a similar metabolic potential (based on completeness-corrected abundances of gene counts) to transform nitrogen and sulfur compounds (PERMANOVA, P = 0.285) (Fig. 5F). In this specific case, the within-group variability was significantly different between both ecosystems (PERMDISP, P = 0.004), demonstrating greater variability in cordgrass-derived MAGs. We also detected some specific trends. For instance, MAGs obtained from cordgrass tend to contain more copies of genes involved in denitrification (e.g., *nap*AB, *nir*BD, *nor*BC), whereas those obtained from the gray mangrove tend to contain more copies of *sox*A and *sox*Z (Fig. 5F). In addition, MAGs obtained from both radicular systems contained up to five copies of *sox*Y, with more variability in cordgrass-derived MAGs (Fig. 5G).

## Discussion

Compartment-specific microbial communities are the pillars of mangrove functioning; therefore, a better understanding of these communities is indispensable to protect, conserve, restore, or rehabilitate this ecosystem (Allard et al. 2020). The phyllosphere of *A. marina* represents a halophilic microbial habitat due to the excretion of salt by the leaf glands. This strong environmental filter contributes to shaping the structure and assembly of leaf-associated microbial communities (Fig. 6). Consistent with this assumption, the phylogenetic null modeling identified homogeneous selection as the predominant assembly process across gray mangrove tree compartments. Nevertheless, homogenizing dispersal may also contribute to the assembly of leaf-associated microbial communities through aerial transport, contact among leaves, and seawater exposure (Meyer and Lindow 2025). Moreover, intrinsic plant characteristics, including the accumulation of trace metals, leaf developmental stages, pH, and size of tree (Lan et al. 2024; Smets et al. 2023), may play secondary but still important roles in the assembly of microbial communities inhabiting gray mangrove leaves.

**Figure 6.**
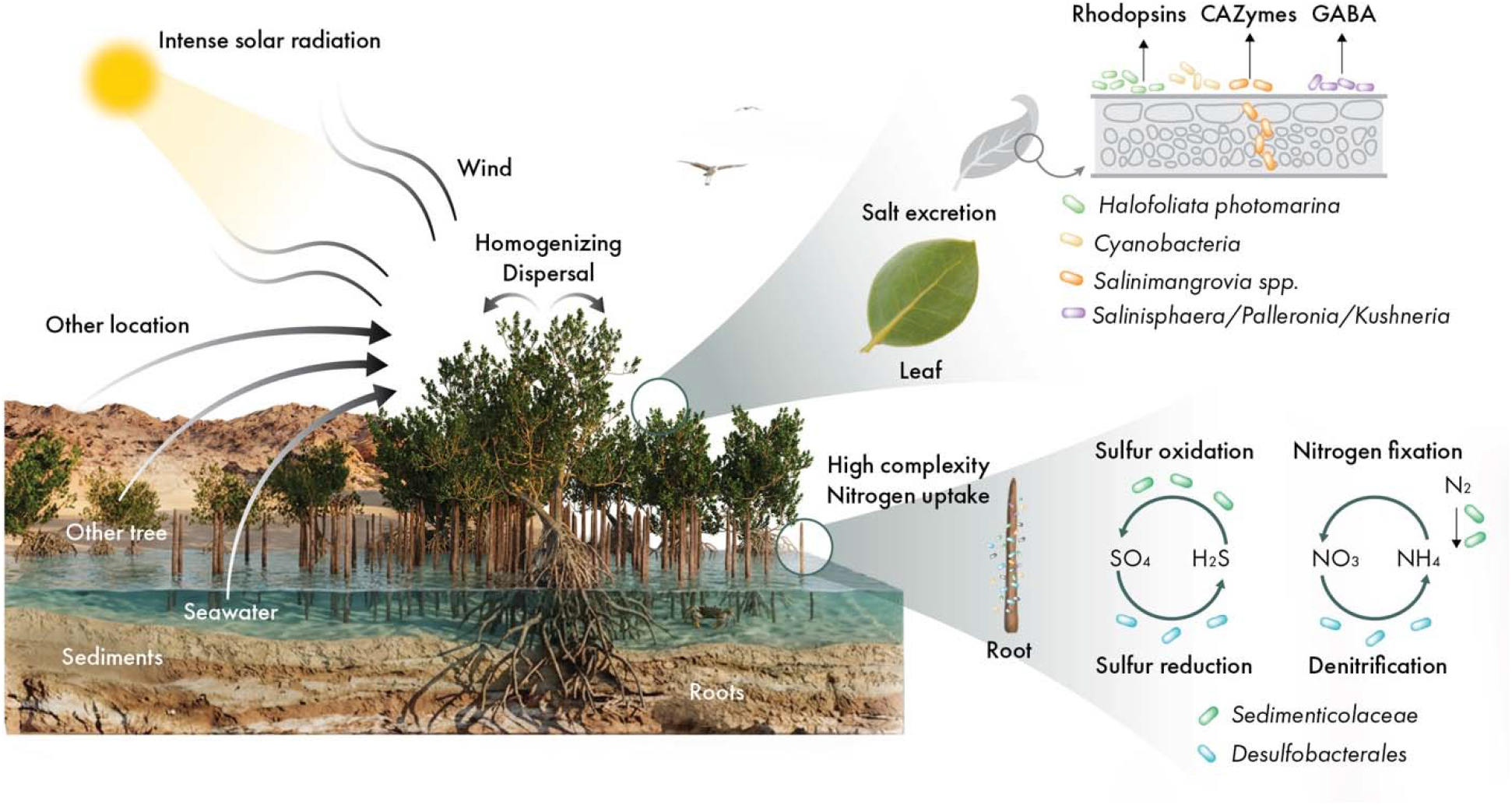
Conceptual figure displaying some of the major findings of our study. This illustration represents two major processes of microbial community assembly in leaves (e.g., selection by salt excretion and dispersal of microbial cells from other locations) and the predictive mechanisms involved in the microbe–mangrove (e.g., γ-aminobutyric acid (GABA) production in leaves), microbe–microbe (e.g., predictive complementary metabolic transformation of sulfur and nitrogen in mangrove roots), and microbe–environment (e.g., phototrophic and ATP generation mediated by rhodopsins in mangrove leaves) interactions. Figure produced by Ana Bigio, scientific illustrator working for KAUST.

Frequent dust storms in the region (Mashat et al. 2020; Alsubhi et al. 2025) could promote the aerial dispersal of microorganisms both among neighboring mangrove trees and across larger geographical distances (Fig. 6). As reported in a salt-excreting desert tree, microbial communities in the phyllosphere are primarily structured by geographical distances rather than plant species (Finkel et al. 2011). The size of gray mangrove trees is also constrained by seawater salinity (Perri et al. 2023), which makes leaves become spatially close to seawater and sediments, thus favoring the dispersion of marine-derived microbial species as is suggested for the *S. alterniflora* phyllosphere (Rolando et al. 2026). In our study, a robust sampling linked to the use of PNA-clamping method allowed the accurate assessment of the diversity and assembly of microbial communities in gray mangrove trees. This provided evidence to confirm that the gray mangrove tree microbiome is organized into distinct microbial signatures across different plant compartments.

The high-salt concentration, intense solar radiation, and predictive poor nutritional conditions turn the gray mangrove phyllosphere into a harsh ecosystem where distinct bacterial and archaeal species can coexist (Yang et al. 2023), probably sharing nutrients and genetic information as mechanisms of survival, adaptation, and evolution. In the present study, two archaeal species of the family *Halococcaceae* were detected in gray mangrove leaves. One represents an abundant novel species (*Halofoliata photomarina* gen. nov. sp. nov.). Although *Halococcus* species have been isolated from the leaf tissue of the black mangrove (*A. germinans*) (Zayas-Rivera et al. 2020), halophilic archaeal populations have been less explored in gray mangrove leaves. The presence of bacteriorhodopsins and halorhodopsins in two MAGs belonged to the proposed novel genus *Halofoliata* suggests that this taxon may be an ancestor of *Halococcus* species, which have lost these genes during evolution (Sharma et al. 2007). This genomic evidence suggests that *Halofoliata* converts solar energy into chemical energy for growth, as an old adaptation to harness and buffer intense solar radiation on gray mangrove leaves (Fig. 6). Furthermore, the bacteriorhodopsin gene found in a MAG affiliated to a *Cyanobacteria* species may represent a signal of an adaptative evolution as a complement of photosynthesis (Hasegawa et al. 2020; Hasegawa-Takano et al. 2024). This putative novel cyanorhodopsin show low similarity compared with *Halofoliata* rhodopsins but contains a domain associated with ketosteroid isomerase–related proteins, which are models for investigating proton-transfer mechanisms (Kędzierski et al. 2020).

A novel bacterial genus identified in this study (*Salinimangrovia arabica* gen. nov. sp. nov) belongs to the order *Rhodothermales* order; abundant taxa being found in the leaves of *A. marina* and *A. germinans* (Alghamdi et al. 2024; Vigneron et al. 2026). As *Salinibacter* species, members of *Salinimangrovia* may possess some physiological similarities compared with extreme halophilic *Archaea* (Mogodin et al. 2005; Oren 2013). Interestingly, *Salinimangrovia* species have the genetic potential to break down complex plant polysaccharides, suggesting that they penetrate plant cell walls and/or generate plant-derived oligo/monosaccharides (glucose, xylose, and arabinose) for their own use or to feed other microbial species (Fig. 6), as reported in *Arabidopsis thaliana* leaf-colonizing bacteria (Hemmerle et al. 2022). Furthermore, *Salinimangrovia* species may be relevant during organic matter turnover, launching decay processes upon senescence. Moreover, MAG-based inferences of physiological traits allowed us to predict that *Salinimangrovia* spp. use D-xylose and L-arabinose as carbon sources for their growth, supporting that they may utilize xyloglucan-derived monosaccharides derived from leaves. These predictive metabolic traits could be useful for the target isolation of this species, as suggested and tested for the cultivation of novel taxa discovered through genome-resolved metagenomics (Karnachuk et al. 2021; Nou et al. 2024; Jiménez et al. 2026).

Although researchers have hypothesized that microorganisms inhabiting gray mangrove leaves can induce plant growth under high-salinity conditions (Alghamdi et al. 2024; Yang et al. 2024), the absence of comprehensive meta-omic analyses has impaired the detection of microbial mechanisms that can improve the tolerance of mangrove trees to stress conditions. We identified that some abundant and widely distributed microbial species found in the leaves of gray mangrove (e.g., *Salinisphaera*, *Kushneria,* and *Palleronia*) (Yang et al. 2023; Alghamdi et al. 2024) have the potential to produce GABA (Fig. 6), a phytohormone that improves osmotic adjustment and photosynthesis efficiency and protects the plant against reactive oxygen species under salinity conditions (Wang et el. 2023; Li et al. 2021; Al-Khayri et al. 2024; Qian et al. 2024; Badr et al. 2024). Interestingly, species from these three genera have been detected, at high abundance, in the cordgrass phyllosphere (Rolando et al. 2026), suggesting that both blue-carbon coastal plants share similar compositional and functional traits in their leaves.

These microorganisms predicted to produce GABA in gray mangrove leaves may utilize putrescine as a precursor. Putrescine is a polyamine that can be accumulated in *A. marina* leaves as a response to high-salinity conditions, acting as an osmo-protectant and antioxidant (Ravi et al. 2020). Apparently, GABA is also accumulated on mangrove leaves, accounting for up to 10%–35% of total free amino acids (Hinokidani et al. 2020); however, a question remains, viz., how much is synthesized by microbes or how much by the plant itself? The microbial-mediated synthesis and accumulation of GABA can be an underexplored extrinsic mechanism used by mangrove trees to increase persistence and resilience to extreme environmental stress and climate fluctuations. In this regard, a controlled induction of this microbial process may be essential for the successful rehabilitation and/or restoration projects of mangroves. The use of genome-based metabolic models (Schäfer et al. 2023) may help predict whether GABA is produced from putrescine and how to induce its accumulation in mangrove leaves.

Microbial communities inhabiting gray mangrove roots and rhizosphere are highly diverse, complex, and specialized due to the selective environmental conditions where they thrive (e.g., low oxygen, high salinity, presence of roots exudates, and different sulfur and nitrogen compounds) (Alzubaidy et al. 2016; Ghabban et al. 2024; Alghamdi et al. 2024). Our genome-centric analysis indicated that two major functional guilds (*Desulfobacterales* and *Sedimenticolaceae*) were abundant in gray mangrove roots. Species of *Sedimenticolaceae* are considered abundant and key microorganisms in blue-carbon ecosystems, driving nitrogen fixation and sulfur transformations (Schmelz et al. 2026; Rolando et al. 2024). They oxidize sulfur compounds (e.g., sulfide) produced by sulfate-reducing bacteria (e.g., *Desulfobacterales*) under anaerobic conditions, allowing the root system to be detoxified by a chemolithotrophic process that requires oxygen, nitrate, or oxidized metals as terminal electron acceptors (Fig. 6). These predictive intricate metabolic interactions between these two functional guilds have been well documented on the roots of *S. alterniflora* and mangrove sediments (Rolando et al. 2022; Nie et al. 2023); however, they have not yet been considered an important microbial signature in *A. marina* roots.

Species from the *Sedimenticolaceae* family are symbiotic sulfur-oxidizing/nitrogen-fixing microorganisms widely found in salt marsh plants, seagrass roots (Schmelz et al. 2026), and lucinid bivalves (Alcaraz et al. 2024). Nevertheless, there exists limited information regarding their presence, distribution, evolution, and metabolic activity in mangrove ecosystems. A previous study reported that nitrogen fixation and sulfur oxidation are generally performed by chemolithoautotrophic *Campylobacteria* species in mangrove sediments (Wang et al. 2024). Hence, we postulate that these species represent a key node in the microbial connectivity among mollusks, seagrass, and mangroves in the Red Sea. Furthermore, we demonstrated that gray mangrove roots host a specific set of *Sedimenticolaceae*-affiliated species that are apparently absent in the roots of *S. alterniflora* but hold a similar genetic potential to perform the same functions within nitrogen and sulfur metabolism. Considering *sox*Y as a predictive marker of evolution (Sudo et al. 2024), our findings suggest that these species have accumulated different copies of *sox*Y as a mechanism of adaptation to the host and to deal with the harsh-fluctuating environmental conditions in the Red Sea. Moreover, the absence of *sox*C in all *Sedimenticolaceae*-affiliated MAGs suggests that these species are generally restricted to environments with low concentrations of oxygen (Ren et al. 2026). Our results infer that an eco-evolutionary coadaptation is imposed onto *Sedimenticolaceae* species inside of gray mangrove roots in the northern Red Sea.

Overall, this study generates substantial advances and foundational knowledge into the microbial ecology associated with gray mangroves trees in the Red Sea. Moreover, our findings suggest that blue-carbon plants possess similar microbial fingerprinting aboveground and belowground. Nonetheless, this may still be a “tip of the iceberg,” and additional investigations are required to accurately determine the activity, dynamics, and global distributions of taxa (rare and abundant) and to validate the predicted functions/mechanisms disclosed during our study.

## Methods

### Locations of gray mangrove forests in the Red Sea

This study was conducted in the Al Wajh Lagoon (∼2000 km^2^), which is located in the northern Red Sea (Saudi Arabia) and hosts one of the northernmost distributions of mangrove forests, predominantly *A. marina.* A sampling expedition was conducted in April 2021 in six distinct geographic zones within the lagoon (Abu Lahiq, Coastal, Um Rumah, Ghawar, Bream, and Shebara) (Fig. 1A). Briefly, Ghawar and Shebara represent mangroves most in contact with the broader Red Sea. Bream is a large island with larger stands of mangroves and seagrass on the sampled side. Um Rumah is the only location sampled where red and gray mangroves grow together. Abu Lahiq lies closer to shore, and Coastal represents the only site on the mainland. Seawater parameters at each location were sampled using a CTD (Ocean Seven Idronaut 310 OS). Salinity ranged between 40 ppt (in Bream) and 48 ppt (in Coastal), and seawater temperature was 25°C–33°C, with a pH of 8–8.2. At each sampling site, three 10-m^2^ plots were established, except for Bream (n = 17). The total number of trees within each plot was recorded, along with tree height and circumference. At each plot, three *A. marina* trees (n = 51) with comparable morphometrics were randomly selected. The height of the sampled mangrove trees was between 69.2 and 308 cm. Four different types of samples associated with *A. marina* trees (leaves, roots, rhizosphere, and sediments) were obtained as described subsequently.

### Sampling of gray mangrove tree compartments

Up to 10 young and healthy leaves of similar size were collected randomly from the bottom, middle, and top sections around each tree. These 10 leaves, which represented an individual sample, were rinsed with sterile Milli-Q water to remove salt water. Next, four leaves were preserved in liquid nitrogen until DNA extraction. Root and rhizosphere samples were collected using a shovel and pruning shears at a short distance (<50 cm away) from the trunk and sometimes submerged depending on the tides. Roots were gently shaken to remove loosely attached sediment and then placed in a sterile falcon tube. Approximately 60–70 g of roots from each tree represents an individual sample. Roots were not surface-sterilized before DNA extraction. Approximately 60–70 g of rhizosphere per sample was collected from the shaken roots. In addition, sediment samples (200 g/sample) were collected in the vicinity of each selected tree (∼10 cm from the trunk) using hand corers sterilized between each tree sampling at a depth of 0–7 cm. Similarly, representative control soil samples were collected from nearby sandy shores devoid of mangrove vegetation at each sampling location. Approximately two-thirds of each sample was snap-frozen in liquid nitrogen for molecular analysis, whereas the remaining material was stored in plastic bags, placed in a cooler, and later transferred to a −20°C freezer until further processing.

### Seawater samples processing

At each sampling site, seawater samples (∼2 L each) were collected into sterile Nalgene bottles stored on ice. After return to the laboratory, the samples were immediately filtered through individually packaged sterile 0.22-μm filter membranes (47-mm diameter; Millipore) mounted on a six-branch aluminum vacuum filtration manifold. All equipment and materials in contact with seawater samples or filter membranes were decontaminated between sampling events by soaking in a 10% bleach solution for 5 min, followed by three rinses with Milli-Q water and a final rinse with ethanol. After filtration, each filter was carefully transferred into an individual 5-mL sterile cryogenic tube and preserved at −80°C until DNA extraction.

### Total DNA extraction

Seawater filter membranes and leaf and root samples were ground in liquid nitrogen using a sterile mortar and pestle. For DNA extraction, ∼200 mg of plant tissue (leaves and roots), ground filter material (seawater membranes), and mangrove sediment samples were used. All samples were homogenized in a bead tissue lyser with chrome beads for 15 min at maximum frequency. DNA was extracted using Qiagen DNeasy PowerSoil Kit (cat. no.: 12855-50) according to the manufacturer’s protocol, with minor modifications. Briefly, ∼200 mg of sample was homogenized with ceramic beads and lysis buffer, followed by precipitation of non-DNA organic and inorganic material. A high-salt binding solution was added to the supernatant, and the lysate was passed through a silica spin column. After removing protein with a washing buffer, an ethanol wash was performed twice rather than once. DNA was eluted with 10 mM Tris, incubated for 2–5 min, and then centrifuged for 1 min. The quality and quantity of DNA were evaluated using a NanoDrop spectrophotometer and Qubit fluorometer (Thermo Scientific), respectively. The extracted DNA was stored at −80°C until sequencing.

### PNA-clamping coupled with 16S rRNA gene amplicon sequencing

The V4 region of the prokaryotic 16S rRNA gene was amplified using the primers 515F (GTGYCAGCMGCCGCGGTAA) and 806R (GGACTACNVGGGTWTCTAAT). Library preparation and sequencing were performed at Argonne National Laboratory (United States) according to a standard protocol (https://github.com/sgreenwald-anl/SeqCore_Protocols) for the Illumina MiSeq platform (2 × 250 bp reads). Polymerase chain reaction was conducted using New England Biosystems’ LongAmp^®^ Hot Start Taq 2× Master Mix with a final primer concentration of 200 pM. Libraries were pooled equimolarly and sequenced at a final concentration of 6.75 pM with a 10% PhiX control DNA. For leaf and root samples (n = 32), additional libraries were prepared using PNA-clamp (GGCTCAACCCTGGACAG) to silence chloroplast sequences (pPNA-S–Chloroplast rRNA Blocker, PP01-25, PNA Bio, https://pnabio.com/product/ppna-s-chloroplast-rrna-blocker/), according to the following PCR protocol: 95°C for 45 s to denature the DNA, with 35 cycles at 95°C for 15 s, 78°C for 10 s, 60°C for 30 s, and 72°C for 30 s, with a final extension at 72°C for 10 min.

### Processing and analysis of 16S rRNA gene amplicon sequencing data

Raw 16S rRNA sequences obtained from 137 samples were processed using DADA2 (v2.24) in R (v4.3.1). Reads were quality-filtered and trimmed (240 bp for forward and 160 bp for reverse reads; maxEE = 2; truncQ = 2; no ambiguous bases), followed by paired-end merging and removal of sequences outside the expected length range (250–256 bp) and chimeras. ASVs were inferred using the DADA2 denoising algorithm, and taxonomy was assigned using DECIPHER (v2.26.0) against the SILVA SSU rRNA database (v138.2). ASVs assigned to chloroplasts (at class level) and mitochondria (at family level) were removed. Alpha diversity (observed ASV richness and Shannon index) and Bray–Curtis beta diversity were calculated using phyloseq (v1.44.0). Differential abundance of ASVs among treatments was evaluated using DESeq2 (v1.51.6).

### Co-occurrence networks and classification of 16S rRNA ASVs

Co-occurrence networks were inferred using SparCC implemented in Python (Friedman and Alm, 2012). Briefly, filtered ASV abundance tables were used as input, and SparCC correlation matrices were generated independently for each treatment to identify positive and negative associations among ASVs. To account for differences in sample size among compartments, 16 samples were randomly selected from each compartment before network inference. To reduce spurious correlations and minimize differences in dataset size among treatments, only the 1000 most abundant ASVs from each treatment were retained for network construction. These selected ASVs represented >90% of the total sequences in each dataset. Correlations were filtered according to magnitude and statistical significance, retaining only strong and significant associations (P < 0.01; r > 0.7 or r < −0.7). The resulting networks were imported into Gephi (Bastian et al. 2009), which was used for visualization and calculation of topological parameters, including degree, modularity, and betweenness centrality. Furthermore, ASVs were classified as generalists, specialists, or rare taxa using the multinomial species classification method implemented in CLAM (Chazdon et al. 2011). CLAM analyses were conducted pairwise among treatments using a coverage limit of 10, a specialization threshold of 2/3, 20 sampling points, and a Bonferroni-adjusted alpha value of 0.05/20. Pairwise CLAM plots were generated using log10-transformed abundance axes to visualize treatment-specific ASV enrichment patterns.

### Assessment of microbial community assembly processes

To infer the ecological processes underlying the assembly of prokaryotic communities, we used a phylogenetic null model based on βNTI in combination with an abundance-based Raup–Crick null model based on Bray–Curtis dissimilarity (RCbray). Analyses were conducted in R (R Core Team, 2024) using functions from the vegan, ape, phangorn, DECIPHER, and Biostrings packages (Oksanen et al. 2025; Paradis and Schliep, 2019; Schliep, 2011; Wright, 2016; Paès et al. 2023). Briefly, ASV sequences were aligned using DECIPHER, and a phylogenetic tree was inferred from the aligned sequences using phylogenetic tools implemented in R. The resulting tree was used for calculating pairwise beta mean nearest taxon distance (βMNTD) among samples. A phylogenetic null distribution was generated using 999 randomizations by shuffling ASV labels across the tips of the phylogenetic tree. βNTI was then calculated as the standardized difference between observed βMNTD and the mean of the null βMNTD distribution, divided by the standard deviation of the null distribution (Stegen et al. 2013). βNTI values >+2 were interpreted as evidence of variable selection, whereas values <−2 indicated homogeneous selection. Values between −2 and +2 were considered not significantly different from null expectations and were further evaluated using RCbray (Stegen et al. 2013).

Pairwise Bray–Curtis dissimilarities were calculated from the filtered ASV abundance table. A null distribution was generated using 999 randomizations according to the RCbray framework, which compares observed abundance-based community dissimilarities against null expectations (Stegen et al. 2013). For each pairwise comparison, the observed Bray–Curtis dissimilarity was compared with the null distribution to calculate RCbray values scaled from −1 to +1. RCbray values between −0.95 and +0.95 were interpreted as stochastic, drift-dominated, or undominated assembly. Values >+0.95 indicated communities that were more dissimilar than expected by chance, consistent with dispersal limitation or other divergent processes, whereas values <−0.95 indicated communities that were more similar than expected by chance, consistent with homogenizing dispersal or convergent processes (Stegen et al. 2013).

### Metagenome sequencing and read-based analyses

Library preparation and sequencing were performed at Argonne National Laboratory (United States). Libraries were prepared using Takara Bio System’s Apollo library prep kit and robot (https://www.takarabio.com/). Specifically, the PrepX Complete ILMN 32i DNA Library Kit, 96 Samples kit was used. Sequencing was conducted on an Illumina NextSeq 2000 using P3 flow cells in paired-end mode (2 × 150 bp). Three flow cells were used, each generating ∼1.2 billion reads. Samples were multiplexed to achieve an average sequencing depth of ∼30 million read pairs per sample (i.e., ∼60 million reads per sample). Raw paired-end metagenomic reads were quality-controlled using fastp v0.23.2 (Chen et al. 2018) with a minimum Phred quality score of 30 to remove adapter sequences and low-quality reads. Quality-filtered reads were used for taxonomic profiling, metagenome assembly, and genome-resolved analyses. Taxonomic classification of clean reads was accomplished using Kraken2 v2.17.1 (Wood et al. 2019) against the PlusPFP database (release June 26, 2026), comprising RefSeq archaeal, bacterial, viral, plasmid, protozoan, fungal, and plant sequences, together with the human genome and UniVec_Core, followed by an abundance reestimation with Bracken v3.0.1 (Lu et al. 2017).

### Reconstruction and processing of MAGs

Quality-filtered reads obtained from root- and leaf-associated sample were assembled independently using MEGAHIT v1.2.9 (Li et al. 2015). In parallel, clean reads were combined and coassembled separately for roots and leaves. Individual assemblies and compartment-level coassemblies were used for maximizing genome recovery across samples with differing microbial abundances. Individual assemblies were binned using MetaBAT2 v2.15.0 (Kang et al. 2019). Coassemblies were independently binned using MetaBAT2, MaxBin2 v2.2.7 (Wu et al. 2016), and CONCOCT v1.1.0 (Alneberg et al. 2014), and the resulting bins were consolidated and refined using metaWRAP v1.3 (Uritskiy et al. 2018). Genome completeness and contamination were assessed using CheckM v1.2.2 (Parks et al. 2015). MAGs with ≥50% completeness and ≤10% contamination were retained for downstream analyses and are hereafter referred to as quality-filtered MAGs. A total of 283 quality-filtered MAGs, comprising 116 from individual assemblies and 167 from coassemblies, were combined and dereplicated using dRep v3.4.5 (Olm et al. 2017). The MAGs were clustered at 95% ANI, using ≥50% completeness and ≤10% contamination as quality criteria. Dereplication yielded 173 nonredundant representative MAGs, including 153 root- and 20 leaf-associated MAGs. Taxonomy was assigned using GTDB-tk v2.6.1 (Chaumeil et al. 2020) against GTDB release R214. Phylogenomic relationships were inferred using GTDB-tk based on concatenated bacterial (120) and archaeal (53) marker genes. Resulting trees were visualized and annotated using iTOL (Letunic and Bork 2021). The distribution and abundance of the 173 representative MAGs were estimated by mapping quality-filtered reads against the dereplicated MAG catalog. Reads were aligned using Bowtie2 v2.5.1 (Langmead and Salzberg, 2012), and alignments were sorted and converted into BAM format using SAMtools v1.16.1 (Danecek et al. 2021). The abundance of MAGs was quantified using CoverM v0.6.1 with the genome workflow and expressed as RPKMs. The resulting MAG-by-sample abundance matrix was used for downstream comparisons among samples and plant compartments.

### Detection and analysis of novel taxa based on MAGs

Average AAI values were obtained using the MiGA platform (Rodriguez-R et al. 2018) against the TypeMat database (release 2025-08). For leaf-derived MAGs, novel taxa were considered based on the following thresholds: new species <95% ANI with its closest relatives and a new genus <65% AAI with related genera (Konstantinidis et al. 2017). High-quality MAGs were uploaded to the Type (Strain) Genome Server (TYGS) to obtain dDDH values and perform phylogenetic comparisons (based on bacterial 16S rRNA gene sequences) and proteome-based phylogenomic analysis (using GBDP) (Meier-Kolthoff and Göker, 2019). Taxonomic placement of proposed novel MAGs was resolved by inferring phylogenomic trees with the members of the family/order with which selected MAGs were affiliated. For this purpose, a set of representative genomes from the type strains identified through the MiGA platform (TypeMat database) and selected MAGs were used to construct a FastTree phylogenomic tree using a single-copy gene set via the GToTree workflow (Lee, 2019). Genome-based prediction of physiological and phenotypic traits was performed using the porTraits workflow of the metaTraits framework (Podlesny et al. 2026) with default parameters.

### Functional annotation of genes detected in MAGs

To functionally annotate genes derived from the 173 quality-filtered MAGs, three complementary pipelines were used, viz., DRAM v.1.5.0 (Shaffer et al. 2020), eggNOG-mapper v2.1.12 (Cantalapiedra et al. 2021), and PGPg_finder v1.1.0 (Pellegrinetti et al. 2024; see below). DRAM integrates gene annotations from multiple reference databases to characterize genome-level metabolic potential. The DRAM distill workflow was used for summarizing gene-level annotations into metabolic modules and pathways, enabling the comparison of the predicted metabolic capabilities across root- and leaf-associated MAGs. For eggNOG-mapper, the eggNOG v5.0.2 database was used, wherein Prodigal for gene prediction and DIAMOND in the sensitive mode were used for sequence similarity searches. mOrtholog assignments were reported using default parameters, including a minimum e-value threshold of 0.001, with no minimum score or taxonomic scope restrictions. KEGG orthology (KO) annotations were used to generate both presence–absence and completeness-corrected abundance matrices (Eisenhofer et al. 2023). KO abundances were estimated using a fractional counting approach, whereby each coding sequence (CDS) contributed a total count of one, divided equally among all assigned KO annotations (i.e., a CDS annotated with n KO annotations contributed 1/n to each KO annotation). Fractional KO counts were summed for each MAG and corrected for genome completeness by dividing by the estimated completeness (expressed as a proportion). Statistical significance was determined using PERMANOVA, followed by permutational analysis of multivariate dispersions (PERMDISP). All analyses were conducted in R using the vegan package.

### Analysis of specific KO annotations

Based on recent studies (Rolando et al. 2024; Whaley-Martin et al. 2023), we selected a set of genes related to sulfur and nitrogen cycle pathways. A schematic overview of the selected genes and their associated metabolic reactions was constructed based on the KEGG database and the sulfur framework reported by Whaley-Martin et al. (2023). To determine whether CDSs annotated as dsrAB were phylogenetically associated with previously described reductive or oxidative types, CDSs assigned to DsrA and DsrB in the recovered MAGs were extracted and independently combined with the corresponding reference sequences from the DsrAB@SILVA database described by Diao et al. (2023). DsrA and DsrB nucleotide sequences were aligned separately using MAFFT (Katoh et al. 2002) with the L-INS-i strategy. Maximum-likelihood phylogenetic trees were independently reconstructed for DsrA and DsrB using IQ-TREE (Nguyen et al. 2015). The best-fitting nucleotide substitution model was selected using ModelFinder, and branch support was evaluated using 1000 ultrafast bootstrap replicates. Furthermore, leaf-associated MAGs were searched for photorhodopsin-related genes, including K04641, K04642, K04643, K06443, K14594, K00514, and K21927. Completeness-corrected abundances of the corresponding KO identifiers were used for comparing the metabolic potential across MAGs.

### Detection of PGP traits

PGP genes were identified using PGPg_finder (Pellegrinetti et al. 2024) against the PLaBAse database with default parameters, including a minimum amino acid similarity cutoff of 50%. PGPg_finder-normalized output tables were used for comparing functional profiles among MAGs and compartments. Heatmaps were generated in R from normalized PGPg_finder summary tables after row-wise standardization of functional abundance values using z-scores, with values capped to improve visualization. For nitrogen-related functions, normalized Level 3 and Level 4 PGPg_finder tables were used to compare leaf- and root-derived MAGs. Differences between compartments were evaluated using Wilcoxon rank-sum tests, with Benjamini–Hochberg correction applied for multiple testing. Overall nitrogen acquisition was visualized using violin plot, boxplot, and jittered point plots, whereas nitrogen subfunctions were represented using log2 fold change values calculated as roots compared with leaves. GABA-related functions were further investigated using gene-level PGPg_finder outputs, focusing on genes assigned to the biosynthesis, conversion, and transport of GABA. These data were visualized as gene-level heatmaps, category contribution plots, and gene prevalence plots across leaf-associated MAGs. Additional taxon-specific comparisons were performed between root-associated *Sedimenticolaceae* and *Desulfobacterales* MAGs using normalized functional profiles, including heatmaps, standardized mean differences, Wilcoxon tests with FDR correction, and principal component analysis based on scaled functional abundance matrices.

### Comparison of gray mangrove and cordgrass-derived MAGs

For comparative analyses, 198 refined MAGs obtained from the root microbiome of *Spartina alterniflora* were downloaded from NCBI BioProject PRJNA1177211 (Huang and Petersen, 2026). To ensure consistent taxonomic assignments across datasets, all MAGs were reclassified using GTDB-tk v2.6.1 with the GTDB reference database release R226. The 198 cordgrass-derived MAGs and the 153 gray mangrove root-derived MAGs recovered in this study were dereplicated using dRep v3.4.5 (95% ANI, minimum completeness 50%, maximum contamination 10%). Dereplication resulted in a nonredundant dataset of 317 bacterial MAGs. Differences in the overall functional profiles of mangrove- and cordgrass-derived MAGs were evaluated using Jaccard distances based on the presence–absence matrix. A second analysis was conducted on *Sedimenticolaceae*-affiliated MAGs using Bray–Curtis distances calculated from completeness-corrected abundances of sulfur- and nitrogen-related KO annotations. The KO annotations contributing most strongly to the observed functional differentiation were identified using the *envfit* function in the vegan package, with significance evaluated by 9999 permutations. Differences in the abundance of *sox*Y were determined using the nonparametric Wilcoxon rank-sum test.

## Data availability

Raw 16S rRNA gene amplicon and metagenome sequencing data are deposited under the NCBI BioProject PRJNA1518281.

## Code availability

Custom scripts for processing, analysis, and visualization of sequencing data are available at https://github.com/mariafpv/RedSea-Mangroves/

## Acknowledgements

This study was funded by the baseline resources (BAS/1/1096-01-01) provided by KAUST to Alexandre Soares Rosado. Thanks to Intikhab Alam for preprocessing and cleaning the metagenome sequences. Thanks also to the RSG team at KAUST, namely Vera Costa for helping with the permits and logistics, Eva Aylagas who supported the team in the field, and Vijayalaxmi Dasari who helped in the lab. Some of the analyses were supported with baseline resources (BAS/1/1109-01-01) provided by KAUST to Susana Carvalho.

## Authors contributions

DJJ: Framed the aims of this study, led the analyses and interpretation of data and results, conceptualized, designed figures, supervised, investigated, and wrote and edited the manuscript draft.

TJ: Analysis of 16S rRNA gene and metagenome sequencing data, and writing, MFPV: Analysis of metagenomes and recovered MAGs, and writing

HA: Sampling, processing of samples, and editing AB: Sampling and processing of samples

KO: Sampling, processing of samples, writing, and editing

LWM: Analysis of 16S rRNA gene and metagenome sequencing data, writing, review, and editing

SC: Sampling, coordination of sampling activities, contribution to project development, writing, review, and editing.

ASR: Conceptualization, funding acquisition, project leadership, resources, writing, review, and editing.

